# Cerebrovascular pathology mediates associations between hypoxemia during rapid eye movement sleep and medial temporal lobe structure and function in older adults

**DOI:** 10.1101/2024.01.28.577469

**Authors:** Destiny E. Berisha, Batool Rizvi, Miranda G. Chappel-Farley, Nicholas Tustison, Lisa Taylor, Abhishek Dave, Negin S. Sattari, Ivy Y. Chen, Kitty K. Lui, John C. Janecek, David Keator, Ariel B. Neikrug, Ruth M. Benca, Michael A. Yassa, Bryce A. Mander

## Abstract

Obstructive sleep apnea (OSA) is common in older adults and is associated with medial temporal lobe (MTL) degeneration and memory decline in aging and Alzheimer’s disease (AD). However, the underlying mechanisms linking OSA to MTL degeneration and impaired memory remains unclear. By combining magnetic resonance imaging (MRI) assessments of cerebrovascular pathology and MTL structure with clinical polysomnography and assessment of overnight emotional memory retention in older adults at risk for AD, cerebrovascular pathology in fronto-parietal brain regions was shown to statistically mediate the relationship between OSA-related hypoxemia, particularly during rapid eye movement (REM) sleep, and entorhinal cortical thickness. Reduced entorhinal cortical thickness was, in turn, associated with impaired overnight retention in mnemonic discrimination ability across emotional valences for high similarity lures. These findings identify cerebrovascular pathology as a contributing mechanism linking hypoxemia to MTL degeneration and impaired sleep-dependent memory in older adults.

## Introduction

Obstructive sleep apnea (OSA) is a common, though underdiagnosed, sleep disorder^1^ that increases in prevalence with age^2^ and is characterized by recurring episodes of complete or partial upper airway collapse during sleep^3^. OSA has been independently associated with reduced regional white matter integrity^4–13^, altered medial temporal lobe (MTL) morphometry^14–20^, cerebrovascular pathology^21–31^, and increased risk for cognitive decline^32–34^. The relative contributions of distinct OSA features (e.g., sleep architecture changes, hypoxemia severity) to the severity of neuropathological, neurodegenerative, and behavioral consequences remains unclear. However, there is growing evidence that intermittent hypoxia, a defining feature of OSA, may contribute to cerebral small vessel disease (CSVD), which may in turn facilitate neurodegeneration, age-related cognitive decline, and ultimately increased Alzheimer’s disease (AD) risk^35–37^.

In recent years, quantification of white matter hyperintensity (WMH) volumes on T2 fluid-attenuated inversion recovery (FLAIR) images has shed light on the contribution of CSVD to age-related cognitive decline^38^. While the exact etiology of WMH is still debated, potential pathogenic mechanisms include hypoxic events which are thought to cause small vessel ischemic episodes leading to cerebral white matter damage and leukoaraiosis^39,40^. In a study of high-risk patients for stroke divided into a nocturnal hypoxia and non-hypoxia group based on oxygen desaturation index (ODI), the group with nocturnal hypoxia had a significantly higher prevalence of silent cerebral infarcts compared to the non-hypoxia group, suggesting a link between hypoxia severity and CSVD development^41^.

Significant evidence suggests that the MTL, which includes the hippocampus and entorhinal cortex (ERC), is particularly susceptible to hypoxia compared to other brain regions--likely due to its relatively higher metabolic demand^42–45^. In a study of middle-aged OSA patients, Canessa and colleagues found reduced gray matter volume in the left hippocampus, left posterior parietal cortex, and right superior frontal gyrus which were consequently associated with cognitive deficits across memory, attention, and executive function domains^46^. In another study of older adult women, OSA was associated with a higher risk of developing mild cognitive impairment (MCI) or dementia in a 4.7-year follow-up, and this effect was linked to the severity of hypoxia^47^. Separately, CSVD appears to impact MTL integrity and memory performance. One study demonstrated that episodic memory decline in older adults with CSVD was explained by temporal interactions between WMH and hippocampal atrophy^48^. In another study of older adults, global cortical thickness and MTL thickness/volume mediated the relationship between WMH burden and global cognition and memory performance, and this relationship was similar across diagnostic groups (i.e., mild cognitive impairment/Alzheimer’s disease)^49^. Emotional mnemonic content seems to be particularly vulnerable to OSA, with a recent study showing fragmented sleep and reduced REM being associated with reduced recognition memory across valences, though this effect was particularly strong for negative content^50^.

It is possible that in patients with OSA, CSVD burden may be driven by specific clinical features of OSA, further leading to MTL degeneration and decline in episodic memory, though these hypotheses have yet to be formally tested. Addressing these critical questions, this study combined high resolution structural magnetic resonance imaging (MRI), clinical polysomnography (PSG), and a sleep-dependent assessment of memory ability (mnemonic discrimination), which is highly sensitive to age and disease-associated changes in MTL structure and function, in a cohort of older adults^51^. The primary hypothesis was that hypoxemia severity would be associated with greater cerebrovascular pathology as measured by WMH burden and that subsequently, greater WMH burden would be associated with reductions in MTL volume and impaired sleep-dependent memory retention. We further hypothesized that WMH burden would mediate the effects of hypoxemia on MTL degeneration and memory deficits.

## Methods

Forty older adults (mean age 72.3±5.8 years, 26 female, Apnea Hypopnea Index (AHI) =13.4±17.4 [range 0-80]) were recruited from the Biomarker Exploration in Aging, Cognition, and Neurodegeneration (BEACoN) cohort, which aims to identify blood and imaging-based biomarkers for preclinical AD in older adults. Of 117 participants in the BEACoN cohort, 40 were eligible and consented to participate in the sleep study and 37 were included in the final analyses. Eligible participants were to be age 60 and older and minimal to no cognitive complaints. Normal cognition was defined as: Clinical Dementia Rating^52^ of 0, Mini-Mental State Examination^53^ score of 25 or higher, and Functional Assessment Staging Tool (FAST)^54^ Stage 1 or 2. Participants were not actively being treated for a sleep disorder and were free of major neurological and psychiatric disorders as assessed during screening with self-report and medical screening questionnaires. No participants had a history of stroke, one had diabetes stable under treatment, two had a history of heart disease, and 13 participants had been or were currently receiving treatment for high blood pressure (around 35% of the cohort included in analyses). Participant characteristics are described in Table 1. Participants did not travel across time zones with a greater than three-hour shift within the three weeks prior to proposed sleep study visit. Participants reported that they were not taking non-SSRI antidepressants, neuroleptics, chronic anxiolytics, or sedative hypnotics.

**Table 1:** Demographics and sleep characteristics of the sample.

| **Demographics** | **N=37** |
| --- | --- |
| **Age, Mean (SD)** | 72.5 (5.6) |
| **Sex, No. Female** | 23 |
| **Years of Education, Mean (SD)** | 16.6 (2.3) |
| **BMI, Mean (SD)** | 24.8 (5.7) |
| **AHI, Mean (SD)** | 13.8 (18.0) |
| REM AHI, Mean (SD) | 25.1 (19.7) |
| NREM AHI, Mean (SD) | 11.3 (18.6) |
| **ODI, Mean (SD)** | 11.0 (15.9) |
| **Race, No.** |  |
| African American or Black | 1 |
| Asian | 10 |
| White | 23 |
| More than one Race | 2 |
| Unknown/Not Reported | 1 |
| **Ethnicity, No.** |  |
| Hispanic | 2 |
| Non-Hispanic | 35 |
| **Vascular Risk History** |  |
| Heart Disease | 2 |
| Stroke | 0 |
| High Blood Pressure | 13 |
| Diabetes | 1 |
| **Antihypertensive/antihyperlipidemic medication (count)** | 15 |
| Both | 4 |
| Only antihyperlipidemic | 4 |
| Only antihypertensive | 7 |
| Neither | 22 |
| **Sleep Report** |  |
| Total Sleep Time (min), Mean (SD) | 336.2 (63.8) |
| Sleep Efficiency, Mean (SD) | 76.1 (14.1) |
| NREM Stage 1 (%), Mean (SD) | 7.9 (4.0) |
| NREM Stage 2 (%), Mean (SD) | 43.6 (8.5) |
| NREM Stage 3 (%), Mean (SD) | 11.9 (8.0) |
| REM (%), Mean (SD) | 12.7 (6.9) |
| Total Arousals Index, Mean (SD) | 24.0 (21.1) |
| Time in Bed (mins), Mean (SD) | 447.2 (72.8) |
| WASO (%), Mean (SD) | 97.7 (64.4) |
| Time Below 90% OSAT (mins), Mean (SD) | 8.4 (15.0) |
| Min OSAT %, Mean (SD) | 86.5 (5.6) |
| REM Min OSAT %, Mean (SD) | 87.3 (6.0) |
| NREM Min OSAT %, Mean (SD) | 88.1 (4.5) |
| **Sleep Disorder Diagnosis, No.** |  |
| Obstructive Sleep Apnea | 24 |
| Central Sleep Apnea | 2 |
| Periodic Breathing | 1 |
| Periodic Limb Movements | 6 |
| Periodic Limb Movement Disorder | 3 |
| Mild Sleep Disordered Breathing | 2 |
| Possible Obstructive Sleep Apnea | 1 |
| Cardiac Arrhythmia | 1 |
| **White Matter Hyperintensity Burden, Mean (SD), cm3** |  |
| Total | 2.12 (2.12) |
| Frontal | 0.85 (.9) |
| Parietal | 0.45 (0.71) |
| Occipital | 0.18 (0.29) |
| Temporal | 0.05 (0.12) |

All participants underwent an in-lab PSG study during which they performed the emotional version of the mnemonic discrimination task^55,56^ delivered in a sleep-dependent manner, with encoding and immediate testing prior to sleep and delayed testing following sleep (see *Emotional Mnemonic Discrimination Task*). In-lab bedtimes mirrored each participants’ habitual sleep time, which was determined by self-report. As part of the parent study, participants underwent MRI at a distinct time point from the overnight study (1.54±0.97 years), which was used to derive brain structure measures and WMH volumes (see *Magnetic Resonance Imaging and WMH Segmentation*). Of 40 subjects, three were excluded in the final analyses due to high motion artifact in MR image (N=2) and one outlier in WMH volume (z-score ±3, Table 1).

All participants gave written informed consent and study procedures were approved by the Institutional Review Board at the University of California, Irvine.

### Polysomnography

Overnight sleep was recorded using standard overnight clinical PSG including electroencephalography (EEG), electrooculogram (EOG), submental electromyogram (EMG), electrocardiogram (ECG), bilateral tibial EMG, respiratory inductance plethysmography, pulse oximetry and a position sensor recorded using Natus® SleepWorks™ PSG Software (NATUS, Pleasanton, CA). Signals were digitized at 1024 Hz. Sleep staging was performed in 30-second epochs using six EEG channels located at approximately 10-20 locations (F3, F4, C3, C4, O1, O2), which were referenced to contralateral mastoid electrodes. Respiratory and arousal events were marked by sleep technicians and were verified by a board-certified Sleep Medicine physician (author R.M.B.), who also scored sleep following American Academy of Sleep Medicine (AASM) scoring guidelines^57,58^. Sleep abnormalities and diagnoses were extracted from the sleep report created by the Sleep Medicine physician (author R.M.B.). Established sleep architecture variables were computed, including total sleep time (TST; total duration of all sleep stages between lights off and lights on), time in bed (TIB), sleep onset latency (SOL; time to first scored epoch of sleep), sleep efficiency (SE; percent of time in bed spent asleep), wake after sleep onset (WASO; duration of wake time following sleep onset), and percent of TST spent in stages N1, N2, N3, and REM. Clinically-relevant measures of sleep disorders were also quantified, including AHI (number of apnea and hypopnea events per hour), respiratory disturbance index (RDI; number of apnea, hypopnea, and respiratory event-related arousals per hour), total arousal index (number of spontaneous and respiratory-related arousals per hour of sleep), and the periodic leg movements during sleep index (PLMSI; number of PLMs per hour). Mean values across the participant cohort for PSG variables are presented in Table 1. All oxygen desaturation events associated with apneas (central, mixed, and obstructive) were demarked by sleep technicians and verified by a sleep medicine physician (author R.M.B.). The most commonly used metrics of hypoxemia were included in this analysis: minimum nocturnal oxygen saturation (Min SpO_2_), oxygen desaturation index (ODI), percentage of total sleep time with an O_2_ saturation of < 90%^59^. Duration, frequency of ≥4% desaturations, and minimum SpO_2_ were stratified by NREM and REM sleep stages. Hypoxic burden was calculated separately for total sleep, REM, and NREM by multiplying respiratory event-associated desaturation percentage by duration of the event, taking an area-under-the-curve approach^60,61^.

### Magnetic Resonance Imaging

MRI data were acquired on a 3.0 Tesla Siemens Prisma scanner with 32-channel head coils at the Facility for Imaging and Brain Research (FIBRE) at the University of California, Irvine. The following structural scans were acquired: Structural T1-weighted MPRAGE (resolution = 0.8 × 0.8 × 0.8 mm, repetition time = 2300 ms, echo time = 2.38 ms, FOV read = 256 mm, slices = 240, slice orientation = sagittal), T2-weighted fluid-attenuated inversion recovery (FLAIR; resolution = 1.0 × 1.0 × 1.2 mm, repetition time = 4800 ms, echo time = 441 ms, FOV read = 256 mm, slices = 160, inversion time = 1550 ms, slice orientation = sagittal) and T2-Turbo Spin Echo (resolution = 0.4 mm × 0.4 mm × 2.0, repetition time = 5000 ms, echo time = 84 ms, FOV read = 190 mm). Two subjects were excluded from final analyses due to high motion (final N=37).

### White Matter Hyperintensities Segmentation

WMH segmentation and lobar parcellation utilized open-source ANTsX software ecosystem^62^ which includes deep learning applications developed for neuroimaging made available for both Python and R via the ANTsXNet (ANTsPyNet/ANTsRNet) libraries. Parcellation was performed using the Desikan-Killiany-Tourville (DKT) cortical labels^63^. An open-source simplified preprocessing scheme was leveraged (e.g., simple thresholding for brain extraction) and an ensemble (N = 3) of randomly initialized 2-D U-nets to produce the probabilistic output^64^, which permitted a direct porting to the ANTsXNet libraries with the only difference being the substitution of the threshold-based brain extraction with a deep-learning approach^62^. All WMH masks resulting from the automated segmentation procedure were manually edited by an expert rater (author B.R.) for improved accuracy.

Segmentation was followed by lobar parcellation, which involved an automated, deep learning-based DKT labeling protocol for T1-weighted images (methods described previously in Tustison et al., 2021). Individual T1-weighted scans were labeled with the cortical DKT regions, the six-tissue regions (i.e., CSF, gray matter, white matter, deep gray matter, cerebellum, and brain stem), followed by a segmentation network applied to the skull stripped image. Parcellations of the frontal, temporal, parietal, and occipital lobar WMH volumes were derived and used for analyses.

### Structural MRI Analyses

Cortical reconstruction and parcellation for cortical volume and thickness measurement and subcortical segmentation was performed with FreeSurfer v.6.0 (Dale, 1999, Laboratory for Computational Neuroimaging, Athinoula A. Martinos Center for Biomedical Imaging, Charlestown, MA, USA http://surfer.nmr.mgh.harvard.edu/) using standard protocols^65,66^. FreeSurfer uses a combined volume-based and surface-based approach to automatically segment the brain and to calculate volume and average cortical thickness in defined regions of interest on T1-weighted MRI volumes. Preprocessing steps included motion correction, averaging^67^, skull-stripping and co-registering to the Desikan-Killiany Atlas^68^. Bilateral regional brain volumes and cortical thickness were defined in subject space based on their T1-weighted image. Segmentations were visually checked for accuracy and manually edited when necessary. Volumes were corrected for total intracranial volume (ICV). Cortical reconstruction was utilized to obtain whole hippocampal volume, ERC thickness, and total ICV.

### Emotional Mnemonic Discrimination Task

Episodic memory was assessed prior to and following overnight sleep using the emotional version of the Mnemonic Discrimination Task (e-MDT)^55,56,69,70^. The MDT is an established framework to tax neural pattern separation processes reliant on the MTL, particularly the ERC and hippocampus, with the emotional variant (e-MDT) additionally allowing for an investigation of how emotion modulates this core computational ability ^69,71–73^. Choice of an emotional variant of the MDT was motivated by evidence that sleep seems to selectively affect consolidation of emotional content^74^, with rapid-eye movement (REM) sleep playing a special role in this process^50^. Prior to sleep, participants were shown a series of 180 emotionally salient images and asked to rate each image on valence (i.e., positive, negative, or neutral) via button press during the initial encoding phase. An immediate recognition memory test followed, wherein 120 images were presented (split evenly among the images presented during encoding (targets), new images (foils), and images that are similar to those in the incidental encoding phase but are not exactly the same (lures). Lures were divided equally into high similarity and low similarity bins. When each image was presented in the immediate test phase, participants were instructed to indicate if they had seen that image before (‘old’) or if it was novel (‘new’) via button press. Following overnight sleep, participants completed the delayed phase of recognition memory testing, wherein they responded ‘old’ or ‘new’ to another set of 120 images, split evenly among targets, foils, and lures. Task trials were randomized and split across immediate and delayed testing to avoid repeating stimuli during testing. Task performance was measured using the bias-corrected Lure Discrimination Index (LDI) for high-similarity lures to focus on the most sensitive measure of Mnemonic Discrimination ability, particularly for age-related changes^75–77^. Sleep-dependent memory consolidation was quantified as the overnight change in LDI (delayed test-immediate test/immediate test).

### Statistical Analysis

To meet assumptions of normality, WMH were normalized via cube root-transformation. Minimum SpO_2_ values were also transformed into a normally-distributed severity index by *sqrt(max(x+1) - x)*, which is a square root transformation for moderately negative-skewed data, meaning greater values indicate lower SpO_2_ percentage (Supplementary Figure 1). Transformed variables were tested using the Shapiro-Wilk test for normality. Outliers that were > 1.5 interquartile ranges from the 1st/3rd quartile were excluded on all variables where transformations failed. In correlational analyses, Pearson’s correlations were computed to test associations. Significant correlations were followed with adjustments for age, sex, and concurrent use of antihyperlipidemic or antihypertensive medications (hereafter referred to as relevant medications) at time of PSG (Table 1). Multiple comparisons were corrected for using the False Discovery Rate (FDR) approach where applicable^78^. Body-Mass Index (BMI) and time between PSG and MRI (mean 1.54±0.97 years) were not significantly associated with any WMH measures and thus were not included in control analyses. In order to assess if WMH burden (M) mediated the relationship between REM hypoxemia severity (X) and MTL gray matter volumes (Y), a mediation model using model 4 in the PROCESS function V.2.16.1^79^ in IBM SPSS^80^ version 27 was constructed, with the same aforementioned covariates included and a 95% confidence interval implemented to infer significance.

## Results

### Hypoxemia is associated with total white matter hyperintensity (WMH) volume

Global WMH burden was positively associated with AHI (r=0.38, p=0.044 FDR corrected), ODI (r=0.39, p=0.044 FDR corrected), minimum SpO_2_ during a night of sleep (r=0.44, p=0.044 FDR corrected), and sleep time below 90% SpO_2_ (τ=0.26, p=0.047 FDR corrected). Hypoxic burden, calculated by using a relatively novel approach of taking the area-under the desaturation curve which has been shown to offer a holistic view of hypoxemia^59–61,81^, was trending though not significantly associated with global WMH burden after FDR correction (r=0.34, p=0.059). In this cohort, total arousals index was not significantly associated with total WMH volume (r=0.23, p=0.201 FDR corrected), and appeared to be orthogonal to sleep minimum SpO_2_ (r=0.19, p=0.239 FDR corrected).

In subsequent models adjusting for age, sex, and concurrent use of relevant medications, only sleep minimum SpO_2_ and time below 90% remained significant predictors of global WMH burden (respectively: overall model adjusted R^2^=0.36, F(4,32)=6.1, p<0.001, sleep minimum SpO_2_ p=0.049 and overall model adjusted R^2^=0.37, F(4,32)=6.3, p<0.001, time below 90% p=0.038). Global measures of OSA (AHI, ODI) did not remain significant (respectively, AHI: F(4,32)= 5.215, Adj. R^2^: 0.319, p=0.005, AHI p=0.173; and ODI: F(4,32)= 5.287, Adj. R^2^: 0.323, p=0.002, ODI p=0.154). In all of these models, age was also an independent significant predictor of total white matter hyperintensity burden, as expected^82^.

### Hypoxemia during Non-Rapid Eye Movement (NREM) and REM sleep and Global White Matter Hyperintensities

Given that cerebral blood flow and cardiometabolic demands of the brain are relatively higher during REM relative to NREM sleep^83–88^, it is possible that associations between global WMH volume and hypoxemia may be particularly strong when hypoxemia occurs during REM sleep. Following the results from the previous section, the next step was to determine if there were sleep stage-specific effects in the hypoxemia-related variables that were robustly associated with global WMH burden. While time below 90% parcellated by sleep stage was not available in this dataset, minimum SpO_2_ in REM and NREM sleep were calculated.

Hypoxemia severity (measured by minimum SpO_2_) during both NREM (r=0.37, p=0.026 FDR corrected) and REM (r=0.49, p=0.004 FDR corrected) sleep were significantly and positively associated with global WMH volume. Subsequent multiple regression models adjusting for age, sex, AHI, and relevant medications revealed that only REM minimum SpO_2_ remained a significant predictor of global WMH volume (overall model adjusted R^2^=0.40, F(5,31)=7.1, p=0.001, REM min SpO_2_ p=0.046), while NREM minimum SpO_2_ did not (overall model adjusted R^2^=0.31, F(5,31)=4.2, p=0.005, NREM min SpO_2_ p=0.485).

### REM Hypoxemia and Lobar White Matter Hyperintensities

Considering the stage-specific findings in REM, the next step was to determine whether the associations between REM-related hypoxemia and cerebrovascular pathology were regionally specific. To do this, REM minimum SpO_2_ was evaluated with lobar parcellations of WMH volume. REM minimum SpO_2_ was positively associated with both frontal (r=0.461, p=0.012 FDR corrected) and parietal (r=0.446 p=0.012 FDR corrected) WMH volumes but not occipital (r=0.118, p=0.648 FDR corrected) or temporal (r=0.078, p=0.648 FDR corrected) volumes, Figure 1. These relationships remained significant with adjustment for covariates for both frontal (adjusted R^2^=0.37, F(4,32)=6.3, p=0.001, REM min SpO_2_ p=0.042) and parietal (adjusted R^2^=0.38, F(4,32)=4.8, p=0.004, REM min SpO_2_ p=0.016) WMH volumes.

**Figure 1:**
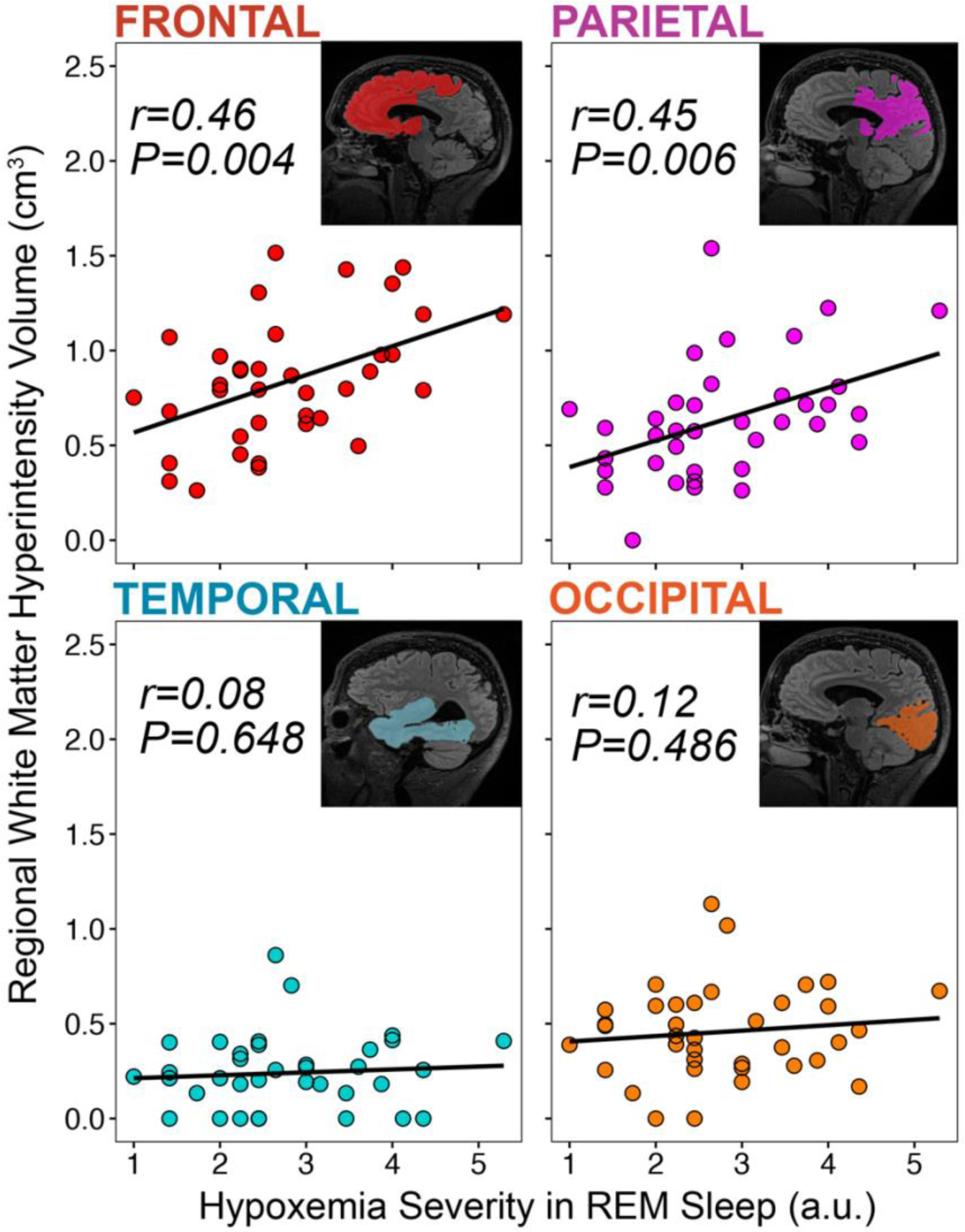
Regional specificity of WMH associations with REM-related hypoxemia. Associations between hypoxemia severity during REM sleep and MRI-measured white matter hyperintensity (WMH) volume in frontal (red), parietal (magenta), temporal (blue), and occipital (orange) cortex. Sagittal slices depicting the region of interest used to assess regional WMH volumes are inset within each respective plot. Hypoxemia severity was computed using a square-root transform of the minimum SpO2 values during REM sleep (sqrt(max(values + 1) - values) due to the negative skew in the across subjects (see Methods for details).

### Frontal White Matter Hyperintensity and Medial Temporal Lobe Volumes/Thicknesses

Since REM-related hypoxemia was related to WMH in the frontal and parietal lobes, these regional WMH volumes were next examined for relationships with MTL structural integrity, which included bilateral hippocampal gray matter volumes and bilateral ERC thickness. ERC thickness was chosen due to the layer-specific degeneration that is observed in aging and AD^89^ and evidence that ERC thinning is a known robust predictor of worse episodic memory performance in aging^90^ and AD^91^.

Greater frontal WMH volume was negatively associated with left ERC thickness (r=-0.44, p=0.028 FDR corrected) and right hippocampal (r=-0.45, p=0.028 FDR corrected) volume (Figure 2). The relationship between left hippocampus was trending but not significant following FDR correction (r=-0.36, p=0.054 FDR corrected) and was not significant with right ERC thickness (τ= −0.051, p=0.657 FDR corrected). Parietal WMH volume was not significantly associated with left (r=-0.31, p=0.104 FDR corrected) or right hippocampal volume (r=-0.38, p=0.054 FDR corrected), though the latter was trending towards significance. Neither left or right ERC thickness was associated with parietal WMH burden (respectively: r=-0.22, p=0.259 and τ=-0.07, p=0.596 all FDR corrected).

**Figure 2:**
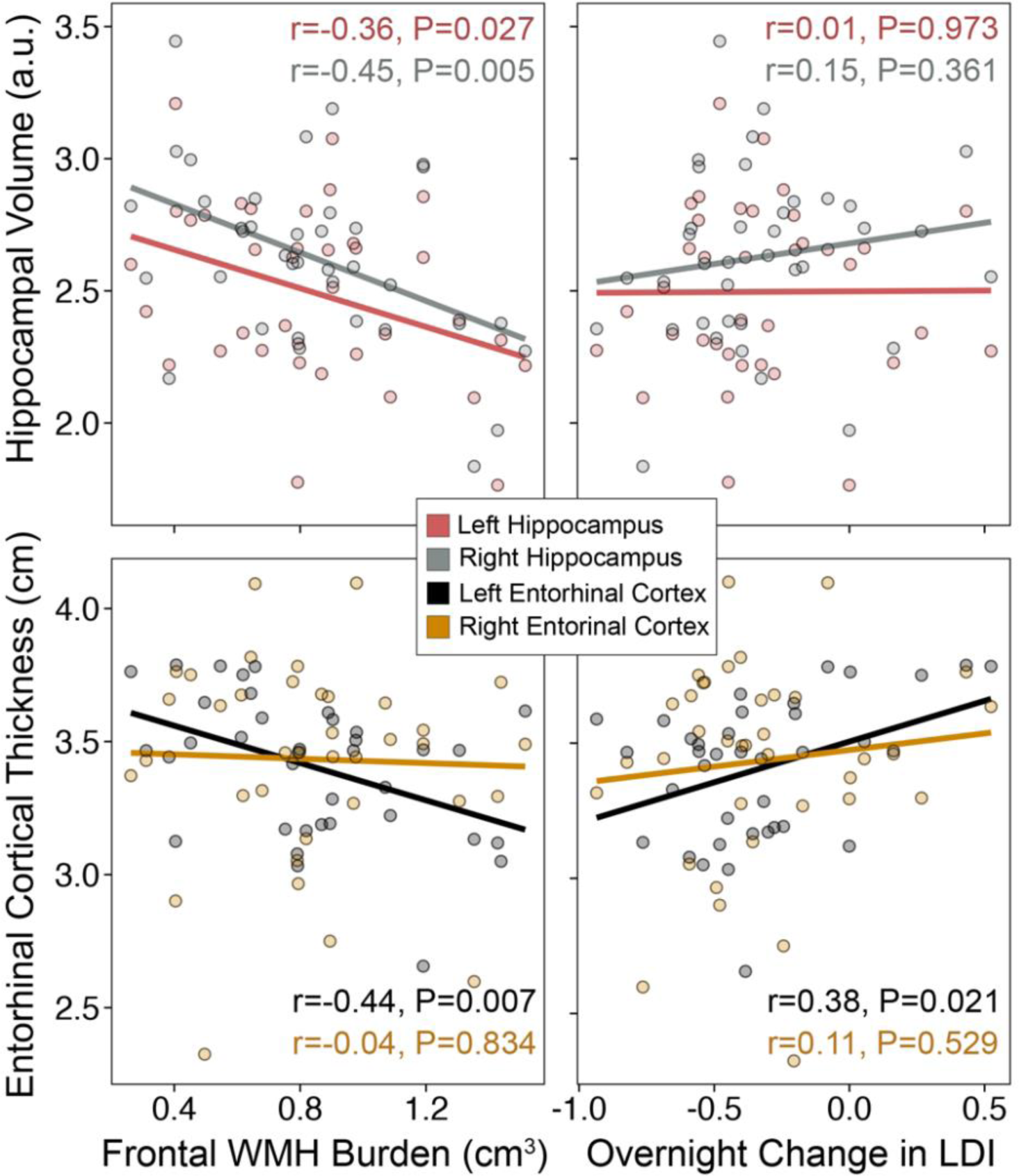
Frontal white matter hyperintensity burden and medial temporal lobe structure and function. Frontal WMH burden associations with bilateral hippocampal volume and bilateral entorhinal cortical thickess, shown in the plots on the left column. On the right, associations between bilateral hippocampal volume and bilateral entorhinal cortical thickess with overnight change in the lure discrimination index (LDI) are shown. LDI was calculated using: p(‘New’|Lure)-p(‘New’|Target). Overnight change was calculated as: overnight change in LDI (delayed test-immediate test/immediate test).

In subsequent models adjusting for age, sex, and use of relevant medications, frontal WMH burden significantly predicted left ERC thickness (adjusted R^2^=0.143, F(4,32)=2.5, p=0.062, frontal WMH volume p=0.008) but not right hippocampal volume (adjusted R^2^=-0.071, F(4,32)=0.405, p=0.803, frontal WMH p=0.582) and not left hippocampal volume (adjusted R^2^=0.185, F(4,32)=3.0, p=0.031, frontal WMH volume p=0.118).

To formally test for an indirect effect of REM-related hypoxemia on ERC thickness through its effects on frontal cerebrovascular pathology, a mediation analysis was performed. Specifically, REM-related hypoxemia severity was the independent variable (X), frontal WMH burden was the mediator variable (M), and left ERC thickness was the dependent variable (Y). While no direct effect was observed (c’=-0.046, p=0.35, Figure 3), a significant indirect effect was observed (indirect=-0.043, 95% CI −0.1174 −0.00015), with the indirect effect accounting for 77.3% of the total effect, Figure 3. Importantly, in a mediation control model, with left ERC thickness as the mediator variable (M) and frontal WMH burden as the dependent variable (Y), no significant indirect effects were detected (indirect=0.0187, 95% CI −0.0247 0.0836).

**Figure 3:**
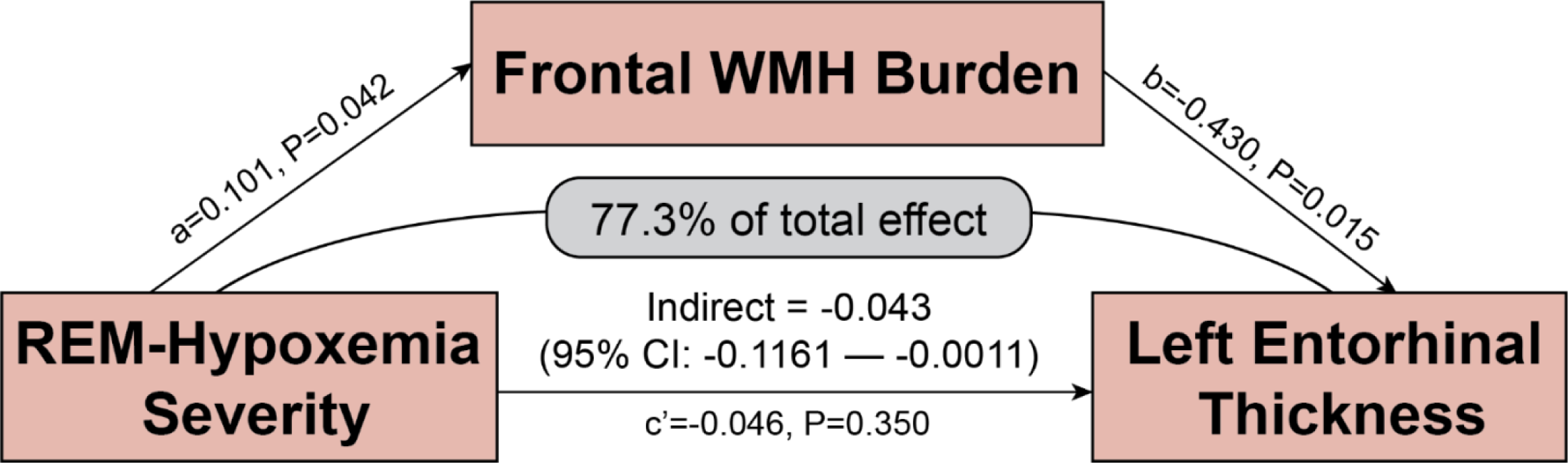
Frontal white matter hyperintensity burden indirectly mediates the relationship between REM-related hypoxia and left entorhinal thickness. A schematic depicting the medial model in which REM-related hypoxemia severity was the independent variable (X), frontal WMH burden was the mediator variable (M), and left ERC thickness was the dependent variable (Y). No direct effect was observed (c’=-0.046, p=0.35). A significant indirect effect was observed (indirect=-0.043, 95% CI −0.1174 −0.00015), with the indirect effect accounting for 77.3% of the total effect. REM-related hypoxemia was positively associated with frontal WMH burden (a=0.101, p=0.042), which in turn was negatively associated with left entorhinal thickness (b=-0.430, p=0.015).

### Entorhinal Thickness and Sleep-Dependent Memory Retention

To examine whether there was a functional consequence of the effects of hypoxemia-related cerebrovascular pathology on MTL-dependent memory function, the overnight change in lure discrimination index (LDI) was quantified for lures across all valences with high similarity to originally presented stimuli as the most sensitive measure of overnight change in sleep-dependent memory retention. Only left ERC thickness was positively associated with overnight change in LDI irrespective of valence (r=0.38, p= 0.021, which remained significant after adjustment for covariates (adjusted R^2^=0.234, F(4,32)=3.7, p=0.013, left ERC thickness p=0.027).

## Discussion

These findings identify cerebrovascular pathology as a candidate mechanistic link between OSA-related hypoxemia severity, MTL degeneration, and sleep-dependent memory impairment in older adults. REM-related hypoxemia, indexed by nadir SpO_2_, may be indirectly associated with impaired overnight MTL-dependent retention of mnemonic discrimination ability due to its relationship with frontal cerebrovascular pathology and associated MTL atrophy.

Furthermore, regional specificity was observed when REM-related hypoxemia was most strongly associated with cerebrovascular pathology in fronto-parietal brain regions. Greater CSVD burden in the frontal region was related to reduced bilateral hippocampal volume and was particularly robust for reduced left ERC thickness, the latter of which was also associated with deficits in sleep-dependent memory. While prior studies have shown that OSA is associated with white matter lesions^21,30^, MTL atrophy^14,20^ and memory impairment^19,50,92,93^, these findings indicate that the effects of OSA are potentially linked with MTL degeneration and memory impairment and are likely downstream of the effects of OSA-related hypoxemia on cerebrovascular pathology. Taken together, these data may partially explain how OSA contributes to cognitive decline associated with aging and Alzheimer’s disease (AD): through cerebrovascular-mediated degeneration of MTL circuits that support memory consolidation during sleep.

The reported association between OSA features and WMH is consistent with prior studies^94–97^. In a large longitudinal study, four unhealthy sleep behaviors including late chronotype, excessive or inadequate sleep duration (>8h or <7h), excessive daytime sleepiness, and snoring (common OSA symptoms), were associated with increased WMH burden^22^. Another group found that moderate OSA severity (AHI ≥ 15), was associated with greater WMH burden, even after adjusting for other risk factors including hypertension^30^. The relationship between OSA and white matter degeneration may be, in part, explained by chronically altered cerebral perfusion in patients with OSA. Studies suggest there is reduced cerebral perfusion during apneic events, which creates an opportunity for ischemic damage particularly at the level of small vessels^98,99^.

OSA is a heterogenous sleep disorder with a complex, multifactorial etiology^100^ which motivates the need to understand how specific features of OSA relate to health outcomes associated with aging and neurodegenerative disease. Given the considerable differences in the neurophysiology supporting NREM and REM sleep stages, understanding stage-specific and sleep feature-specific aspects of OSA would allow for development of more targeted therapeutics and novel clinically-relevant biomarkers of the health consequences of OSA. For these reasons, it is noteworthy that the strongest associations were specifically detected with the severity of hypoxemia during REM sleep. Global measures of OSA severity that collapse across NREM and REM sleep stages may not be sensitive to the neuropathological and cognitive consequences of OSA, since NREM sleep duration dominates total sleep time^83^. Non-specific assessment of OSA could result in underestimation of hypoxic burden pertinent to cognitive health, particularly in patients with a REM-dominant expression of OSA^101,102^. In the Wisconsin Sleep Cohort study, 18% of the sleep studies with no evidence of OSA (i.e. overall AHI <5) demonstrated moderate or severe OSA during REM sleep (i.e. REM AHI ≥15)^102^. A significant percentage of OSA patients (14–37%) have most of their respiratory events during REM sleep^102,103^. In conjunction with the current findings, hypoxemia severity and hypoxic burden have been consistently associated with clinical outcomes including cardiovascular disease related-mortality and incident heart failure^61,81^. In a study of sleep-disordered breathing in older women, elevated ODI was associated with increased risk for developing MCI or dementia, whereas measures of sleep fragmentation and sleep duration were not, supporting results presented here and further implicating hypoxic burden in cognitive deficits and pathogenic mechanisms of cognitive decline^47^. Future studies following individuals longitudinally should examine whether trajectories of neurodegeneration and cognitive decline differ in individuals matched for overall OSA severity but with more REM-dominant versus more NREM-dominant OSA expression patterns in hypoxic burden.

There are several possible explanations as to why the brain may be particularly vulnerable to cerebrovascular injury resulting from OSA-related hypoxemia occurring during REM sleep. First, REM sleep is associated with higher sympathetic activity and cardiovascular instability in healthy individuals and magnified in patients with OSA^86,104^. Patients with OSA exhibit increased sympathetic tone, both throughout the night and during the day^104,105^. There is also a reported increased tendency for upper airway collapse during REM sleep due to decreased genioglossus muscle tone^102,106^, which may explain why obstructive apneas and hypopneas in REM are longer in duration and are associated with significantly greater oxygen desaturation when compared to events occurring during NREM sleep^107,108^. Indeed, ventilation is more erratic and SpO_2_ is less well maintained during REM than NREM sleep^108^, with approximately a twofold increase in variability in respiratory rate during REM^109,110^. Other studies have found greater reduction in ventilatory response to nocturnal hypoxemia during REM than NREM sleep^111^. In REM sleep, baseline SpO_2_ can be lower, lending to a larger precipitous fall in SpO_2_ with apnea occurrence^112^. Steeper desaturations or a lower baseline SpO_2_ during REM sleep may also be explained by evidence that lung volume may be lower during REM sleep^113^, making the pulmonary oxygen reservoir lower and creating an environment in which SpO_2_ is likely to fall faster. In comparative metabolic demand, higher oxygen consumption during REM than NREM sleep could lower SpO_2_ levels in REM^112^. Taken together, these acute changes in hemodynamics and ventilatory control during REM sleep in patients with OSA could play a part in triggering ischemic events^102,111,112,114^. With regard to cerebral perfusion, fluctuations appear to be greater for patients with OSA during REM sleep transitions than during NREM sleep transitions^97^. Early studies in healthy adults have supported the idea that cerebral blood flow seems to be generally reduced during NREM sleep and an increased during REM sleep as compared to wake^84,85,115–119^. Positron emission tomography studies have also shown widespread brain activation in REM compared to NREM, similar to wake^120–122^. Given that these findings might reflect greater cerebral metabolic demand during REM compared to NREM sleep, oxygen desaturation occurring in REM sleep may be more likely to lead to ischemic events in the brain when oxygen requirements are high and may render regions with relatively higher metabolic demand such as the hippocampus at greater risk of hypoxic injury. Though there was no significant difference between minimum SpO_2_ in REM versus NREM in this sample, differences in physiological environment (ventilatory drive, lung O_2_ reserve, metabolic demand) between NREM and REM sleep may confer differential impacts of nocturnal hypoxemia to the brain in a stage-dependent manner. Taken together, these data support the hypothesis that respiratory events in REM sleep may play a larger role in triggering gradual loss of adaptive physiology underlying cerebrovascular architecture to cope with ischemic events over time compared to events in NREM sleep^8^.

The findings presented here also demonstrate regional specificity to CSVD burden associated with REM-related hypoxemia. Fronto-parietal lobes have been consistently referenced in relation to OSA and cognitive functioning deficits with sleep loss or fragmentation shown to be associated with fronto-parietal atrophy^123^. Consistent with this sample, Kim et al. found that WMHs were predominantly seen in the frontal lobe of middle-aged to older adults^30^. In a study of patients with severe sleep apnea evaluated, authors found N-acetylaspartate-to-creatine and choline-to-creatine ratios were significantly lower in the frontal white matter of OSA patients compared to controls^124^, which may be representative of impaired energy metabolism and neuronal injury of various causes^125^. In another study, more respiratory events during REM sleep were associated with reduced daytime regional cerebral blood flow (rCBF) in the bilateral ventromedial prefrontal cortex and in the right insula extending to the frontal cortex^101^, and similar results were found in another study where CBF was significantly reduced during sleep in frontal and occipital cortex, pons, and the cerebellum in patients with OSA relative to controls^97^.

The hippocampus and the ERC structures of the MTL show early histological changes in Alzheimer’s disease^126,127^. The ERC is known to act as the major input-output structure of the hippocampal formation and nodal point in cortico-hippocampal circuits, stabilizing mnemonic content for long-term storage^128^. Consistent with the presented study findings here, these two regions of the MTL, the hippocampus and ERC, have been shown to be impacted by OSA and hypoxia more broadly in prior studies^6,14,17,20,35,129–138^. These effects are particularly apparent in older adults with subjective and objective cognitive difficulties, where ODI was associated with reduced cortical thickness in the bilateral temporal lobes which in turn was associated with reduced verbal encoding^139^. These studies have identified both altered brain metabolism and neuronal loss in the hippocampus and frontal cortex in patients with OSA^6,140,141^. Rodent studies mirror these findings. One stereological study of hippocampal subregions revealed that both white and gray matter in CA1 and CA3/DG subfields were particularly vulnerable to hypoxic injury^35^, while another showed that hypoxia-related decreases in CA1 and DG volumes were due to neuronal cell death-mediated brain atrophy^132–134^. These findings have major implications in pattern separation processes, which are supported by the hippocampus^142^ and may explain sleep-dependent memory deficits in people with OSA. However, it has remained unclear how OSA-related hypoxemia results in MTL degeneration. The findings presented identify cerebrovascular pathology as a candidate mechanism driving MTL degeneration associated with OSA-related hypoxemia. Prior studies also support this possibility, suggesting a strong link between WMH burden and MTL atrophy^143–146^ which the present findings extend by implicating hypoxic burden during REM sleep as a contributor to this link. AD patients with periventricular WMH had a significantly increased risk of progression of MTL atrophy compared to those without baseline periventricular WMH^144^. Findings presented here add a layer to this story, indicating that OSA-related hypoxemia severity during REM sleep is associated with MTL degeneration through its association with frontal cerebrovascular pathology. Thus, these neuropathological consequences of OSA are not independent but potentially linked with each other and can explain the cognitive consequences typically seen in OSA.

This study found that lower left ERC thickness was associated with greater overnight reductions in mnemonic discrimination ability for highly similar stimuli across all valences. Prior studies have shown age-related deficits in object pattern separation associated with hypoactivity of the anterolateral ERC in cognitively asymptomatic older adults^147^, and another study showed thickness in the ERC and subicular cortices of MCI subjects at initial assessment correlated with changes in memory encoding^148^. The presented findings here are consistent with studies showing memory impairments, including for sleep-dependent memory, in patients with OSA^50,92,93,149^. While prior work has separately identified that OSA impacts MTL structure and sleep-dependent memory impairment, the results of the current study extend this literature, linking OSA to sleep-dependent impairment in mnemonic discrimination ability in older adults through downstream consequences of OSA on MTL structural integrity, as would be predicted by contemporary models of the role of the MTL in pattern separation and sleep-dependent memory^73,142,150^. Previously, Cunningham and colleagues reported that reduced REM sleep was associated with disruptions in overall recognition and veridical memory for emotional content^50^, which may explain why greater hypoxemia severity in REM sleep might impact overnight memory consolidation. This study has several limitations. First, the relationships are correlational and the contribution of independent factors also prevalent in OSA, such as hypertension, atherosclerosis, metabolic syndrome, and atrial fibrillation^97^ are not fully characterized in this cohort. However, the current study adjusted for concurrent use of antihyperlipidemic and antihypertensive medications to minimize these confounds. Importantly, in this cohort, no participants had a history of stroke, one had diabetes stable under treatment, two had a history of heart disease, and 13 participants had been or were currently receiving treatment for high blood pressure (around 35% of the cohort included in analyses). Regardless, this issue remains an important concern, given prior studies linking OSA severity during REM sleep with hypertension^102^. Hypertension is common in older age and a major CVD risk factor^151^, indicating that comorbid OSA and vascular risk from other sources may confer even more risk for WMH-related neurodegeneration and cognitive deficits. Second, this study sample was not statistically representative of the racial-ethnic makeup of the United States, which limits generalizability. This is important to highlight, as Black/African American adults are 88% more likely to have OSA and more than twice as likely to have severe OSA compared to their non-Hispanic White counterparts^152,153^ and also have an increased risk for stroke^154^ and other cerebrovascular disease^155^. Third, this study was not powered to examine sex differences in relationships among OSA severity, WMH volumes, and MTL structure and function. Prior studies show that women may express a larger portion of their OSA events during REM sleep relative to men^103,156^, and there are sex differences in the relationship between OSA and both white matter integrity and MTL regional volumes^129,157^. Future studies should explore if REM-dominant OSA expression may be a particularly significant contributing factor to sex differences in MTL atrophy and cognitive decline in aging and AD.

## Conclusions

In conclusion, OSA-related hypoxemia was associated with fronto-parietal cerebrovascular pathology, which was associated with MTL integrity and sleep-dependent memory consolidation. Provided that hypoxemia severity during REM sleep was the strongest predictor of WMH volume indicates that even those with lower global OSA severity but higher levels of hypoxemia during REM sleep may still be at risk for neuropathologic change and related memory impairment. This finding underscores the necessity to examine the cognitive benefits of OSA treatment in patients with moderate to severe REM-related hypoxemia even if global OSA severity measures remain low. Future studies following individuals longitudinally should examine whether trajectories of neurodegeneration and cognitive decline differ in individuals matched for overall OSA severity but with more REM-dominant versus more NREM-dominant OSA expression patterns, as these identified pathological changes may be contributing mechanisms linking OSA to increased Alzheimer’s disease risk^33,36,158^.

## Funding information

National Institute on Aging K01AG068353

National Institute on Aging R01AG053555

National Institute on Aging F31AG074703

National Institute on Aging R21AG079552

American Academy of Sleep Medicine Foundation SRA-1818

---

**Supp. Figure 1:**
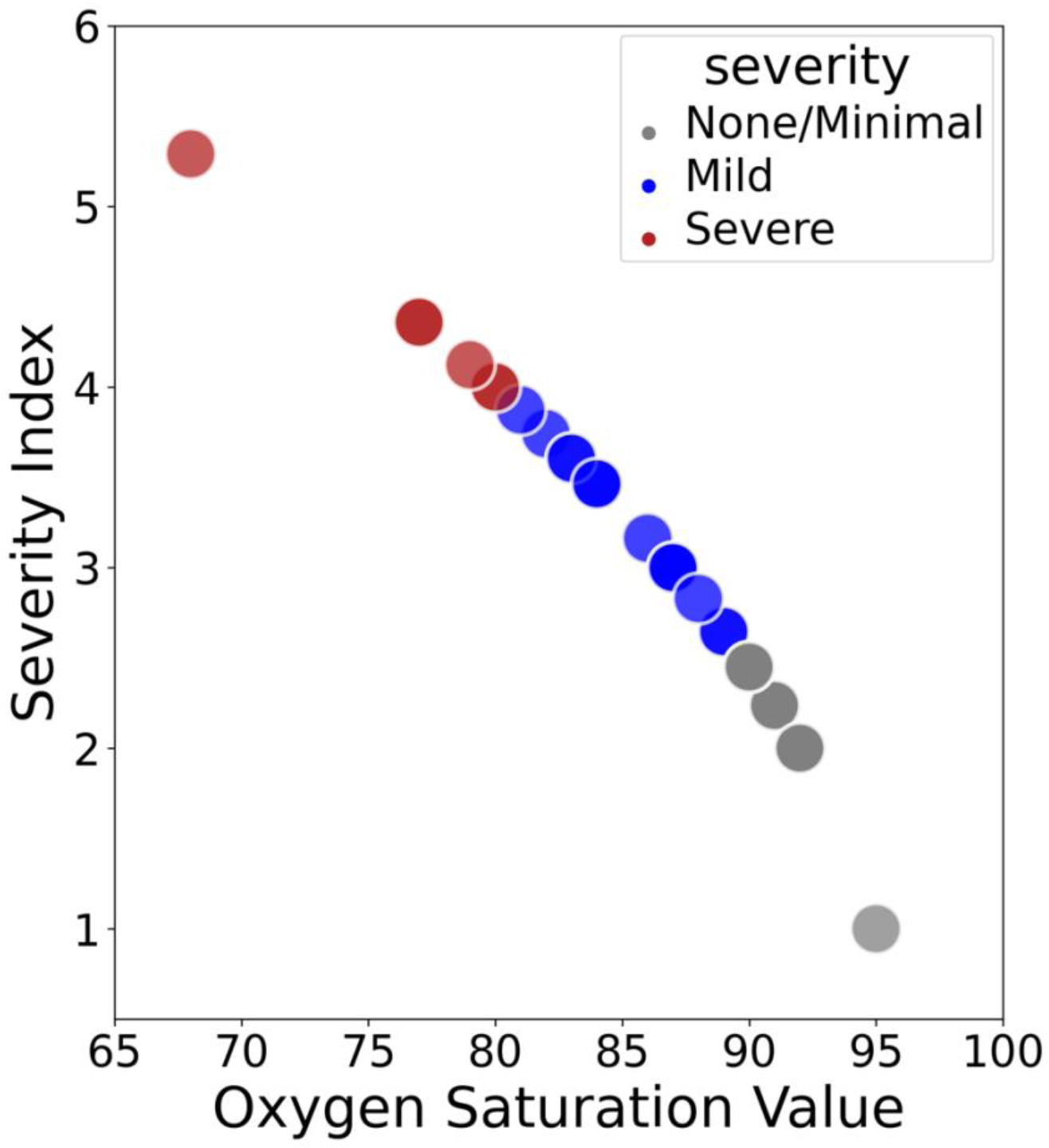
Depicts the minimum SpO2 transformation into a normally-distributed severity index using sqrt(max(x+1) - x), a type of square root transformation for moderately negative-skewed data, meaning greater values indicate lower SpO2 percentage.

